# Maximization of Energy Recovery from Starch Processing Wastewater by Thermophilic Dark Fermentation Coupled with Microbial fuel Cell Technology

**DOI:** 10.1101/2022.10.25.513786

**Authors:** Mohit Kumar, Soumya Pandit, Vinay Patel, Debabrata Das

## Abstract

Utilization of organic wastewater for hydrogen production has dual advantages of clean energy generation and bioremediation which is sustainable for a longer period. To maximize the energy recovery from starch rich wastewater, a two stage system comprising of thermophilic dark fermentation coupled with microbial fuel cell was employed. A single parameter optimization strategy was implemented for the operation of the batch system. The maximum cumulative hydrogen production obtained was 2.56 L L^−1^ with a 48 % reduction in COD under the optimal conditions of 35 g L^−1^ initial substrate concentration (COD), temperature 60 ^o^C, and pH 6.5. The H_2_ yield and H_2_ production rate were 6.8 mol H_2_/kg COD_reduced_ and 731.3 mL L^−1^ h^−1^ respectively. The effect of the organic loading rate (OLR) on H_2_ production rate was studied in a continuous stirred tank reactor (CSTR). A maximum hydrogen production rate of 913 mL L^−1^ h^−1^ was observed at an OLR of 5.6 g L^− 1^ h^−1^. Effluent recycle played an important role in the improvement of H_2_ production. A maximum H_2_ production rate of 1224 mL L^−1^ h^−1^ was observed at a recycle ratio of 0.6. Power density of 4.2 W m^−3^ was observed with MFC using the dark fermentative spent media neutralized with carbonate buffer at an optimal pH of 7. A total COD reduction of 86% was observed.

## 1. Introduction

Production of clean energy and utilization of wastewater for bioenergy production is a novel and promising approach to meet the increasing energy needs as a substitute for fossil fuels. H_2_ is a clean fuel with no CO_2_ emissions and can easily be used in fuel cells for the generation of electricity. Among the different gaseous fuels, H_2_ has the highest calorific value of 3042 cal m^−3^ (considering water as a product) [1]. Dark fermentation and photo-fermentation are the two possible ways of H_2_ production from organic wastes. Dark fermentation is preferred over the photo-fermentation because of its higher production rates and the ability to grow in the absence of light. The dark fermentative H_2_ production using organic wastes is an attractive option of waste treatment as well as sustainable energy production. Dark fermentation at temperatures in the thermophilic range has a considerable advantage over the mesophilic temperatures because of favorable thermodynamics at elevated temperatures and less susceptibility to methanogenic contamination.

Organic pollutants are anaerobically converted into various volatile fatty acids (VFA) like acetate and butyrate in fermentation; in this process, H_2_ is a byproduct. The major bottleneck associated with the dark fermentative biohydrogen production process is the partial substrate utilization [2]. As a result, a high amount of unutilized substrate along with VFA generated during dark fermentation usually remain in the effluent. These soluble metabolites can be converted into methane by the action of methanogens or can be utilized by photo fermentative bacteria for hydrogen production or the production of electricity by the action of exoelectrogens in microbial fuel cells [3,4]. While methanogenesis is a kinetically slow process, the high opacity of the spent-medium renders it unsuitable for high yield photofermentation. Recently, the microbial fuel cell is emerging as an attractive option for treating the dark fermentative effluent for further COD reduction and electricity generation[5].

In MFC, electrons involved in the metabolic pathways are redirected for the production of electricity. MFC is receiving renewed attention because of the intriguing possibility of developing sustainable energy processes that do not require the consumption of fossil fuels. So far, the current and power densities achieved with MFCs are relatively low but still could be used to power sensors and other electronic devices, and treat waste while generating electricity [6].

In the present study, efforts have been made to recover bioenergy through a two-stage dark-fermentation followed by MFC using starch processing wastewater to make the process eco-friendly and economical. Batch fermentation with starch processing wastewater as substrate was carried out. Furthermore, continuous hydrogen production was performed in a chemostat and the effect of organic loading rate on H_2_ production was studied. The effect of effluent recycle on the rate of hydrogen production was also studied. The spent medium of the dark fermentative H_2_ production process was used as anolyte in a single chambered MFC (sMFC). The performance of sMFC was evaluated based on the volumetric generation and wastewater treatment efficiency in terms of coulombic efficiency (CE). Overall energy efficiency, COD, and carbohydrate removal efficiency were evaluated to show the efficacy of the whole integrated process.

## 2. Materials and methods

### 2.1 Microorganism and culture conditions for dark fermentation

Sludge was collected from an anaerobic digester. A suitable consortium capable of H_2_ production was developed and used in all the experiments [7]. The consortia was regularly maintained anaerobically in serum bottles by subcultures. The medium used for maintenance comprised of, glucose (10 g L^−1^) yeast extract (4 g L^−1^), tryptone (10 g L^−1^), FeSO_4_ (20 mM), and Cysteine HCl (1 g L^−1^) (DSMZ medium of No141, German Collection of Microorganisms and Cell Cultures) [7].

Starchy processing wastewater was collected from local households in Kharagpur, West Bengal, India. The medium used for production contained different dilutions of the wastewater as the carbon source in place of glucose while the remaining media components remained the same as mentioned above.

### 2.2 Single parameter optimization

Earlier studies have shown that hydrogen production is a function of limiting substrate concentration, pH, and temperature [8]. Single parameter optimization was done with respect to initial substrate concentration (COD), pH and temperature by varying one parameter, at a time, while keeping the other two parameters constant. This study was carried out in 100 mL serum bottles with 70 mL working volume under strictly anaerobic conditions maintained by purging with nitrogen for 5 min.

The activation energy for H_2_ production was calculated using the Arrhenius equation, given by,

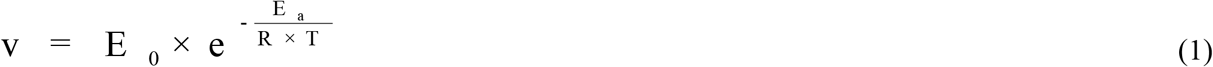

Taking natural logarithm on both the sides, we get

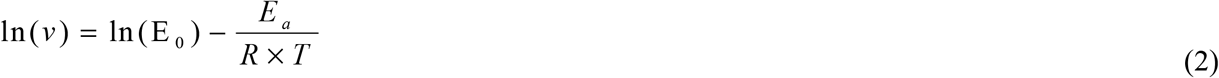

By plotting the graph of ln(v) vs 1/temperature the activation energy of the mixed culture was determined. Where v is the rate of H_2_ production (mmol L^−1^ h^−1^), E_o_= Arrhenius constant, E_a_ = Activation energy (KJ mol^−1^), R= gas constant (8.314 KJ Kmol^−1^ K^−1^) and T is the temperature (^o^K).

### 2.3 Batch fermentation under optimized conditions

Medium containing the starchy processing wastewater was subjected to batch fermentation under optimized conditions of temperature, pH, and initial substrate concentration values obtained from single parameter optimization studies. Batch fermentation was carried out in 500 mL customized double jacketed reactors with 400 mL working volume. Purging with nitrogen was carried out to maintain anaerobic conditions [9].

### 2.4 Continuous production of hydrogen in continuous stirred tank reactor (CSTR) Experimental setup

Continuous hydrogen production was carried out in 500 mL customized double jacketed continuous stirred tank reactor with a working volume of 400 mL. The reactor contents were mixed by a magnetic stirrer. Temperature control was achieved by using a circulating water bath maintained at 60 ^°^C. Two peristaltic pumps were provided at inlet and outlet lines to maintain the desired flow rates. A recycle line was also provided to recycle a part of the effluent stream. A feed tank of 10 L was provided for the storage of medium for fermentation. The gas generated during the reaction was collected by a water displacement system through an outlet provided at the top of the reactor [10]. A quasi-steady state (5% variation) was observed at each dilution rate with respect to constant values of H_2_ evolution rate, glucose, and cell mass concentration in the effluent. The experiments were repeated at different flow rates to get maximum hydrogen production and sugar utilization.

### 2.5 Anolyte and Inoculum for MFC

The acid-rich effluent generated after thermophilic dark fermentative H_2_ production using starchy wastewater was subsequently used as a substrate for bioelectricity generation using a single chambered MFC. Two sMFC set-ups were developed, in duplicates, referred to as sMFC-1 and sMFC-2. The chemical oxygen demand (COD) of the spent medium was diluted and kept in the range of 2990 to 3010 mg L^−1^ for both the MFCs. The initial pH in one of the sMFCs was maintained in the range of 7.0 by the addition of suitable alkali (sMFC-1), while in other one (sMFC-2) was kept undiluted while maintaining same pH with alkali addition. Anaerobic mixed consortia obtained from the bottom sludge of the septic tank of IIT Kharagpur was used as parent inoculum. The inoculum sludge was sieved through 1-mm sieve, preheated at 100 °C for 15 min and was allowed to cool. Before that, the culture was washed thrice in a saline buffer, with intermittent centrifugation at 5000 rpm, and enriched in synthetic wastewater, under an anaerobic microenvironment at pH 7 on a shaker set at 100 rpm at room temperature. The resulting enriched culture was inoculated along with the feed.

Initially, during a week long adaptation period, the MFCs were operated in an open circuitry configuration in fed-batch mode at ambient temperature (29 ± 2°C) using sucrose based synthetic water as per the composition provided in the reference [11]. Subsequently, after achieving stabilized performance, after a week, the reactors were operated on the spent medium of dark fermentation. Prior to this, the spent medium after dark fermentation was centrifuged at 5000 rpm (Sigma Sigma 3K30), the supernatant was collected and supplemented with carbonate buffer (MFC-1) or alkali solution (MFC-2) for anolyte preparation. To encourage microbial growth, in both the sMFCs, the pH of the wastewater was maintained near neutral using carbonate buffer (Na_2_CO_3_ and NaHCO_3_) and alkali (NaOH) solution. A third sMFC reactor was set-up, in duplicate, to serve as control (sMFC-3) where the initial pH of the anolyte remained unaltered. Subsequent to stabilization, the MFCs were operated in close circuit configuration fed with spent media of dark fermentation, after dilution in fed-batch mode, for 9 consecutive cycles (36 h cycle^−1^) at ambient temperature (29 ± 2°C) and pressure.

### 2.6 MFC configuration

Six identical cuboidal-type sMFC were developed for the experiments. Each reactor was of 100 mL capacity (working anolyte volume 0.1 L) with a side opening where the membrane cathode assembly was connected. In order to prepare a membrane cathode assembly (MCA), the cathode was made of carbon cloth (projected surface area = 16 cm^2^). Nafion 117 (DuPont, Wilmington, DE) was hot pressed directly onto the catalyst [MnO_2_-NT] and ultrasonically mixed carbon black Vulcan XC-72; [MnO_2_-NT loading = 0.1 mg cm^−2^; Vulcan XC-72=3mg cm^−2^] loaded side of carbon cloth by heating it to 125°C at 1780 kPa for 3 min [12]. The MFC consisted of an anode and cathode placed on opposite sides. The non catalyzed ammonium pre-treated carbon cloth was used as anode (2×3 cm^2^). sMFCs were operated in fed-batch mode at a fixed external resistance of 100 Ω (except as indicated) and refilled after 36 h.

### 2.7 Analytical methods

Total solid (TS), volatile solid (VS), COD, and alkalinity were determined using standard techniques [13]. H_2_ content in the evolved gas was analyzed using a gas chromatograph (GC; PerkinElmer LLC, MA, USA) equipped with a thermal conductivity detector (TCD) and a stainless-steel column packed with Porapak Q (80/100 mesh). The operational temperatures of the injection port, oven, and detector were 80 °C, 150 °C and 200 °C respectively. Volatile fatty acid (VFA) analysis was done in gas chromatography (PerkinElmer LLC, MA, USA) equipped with flame ionization detector (FID) and stainless-steel capillary column coated with 10 % PEG-20 M and 2 % H_3_PO4 (80/100 mesh). Injection port, detector, and programmed column were maintained at 220 °C, 240 °C, 130–175 °C, respectively. H_2_ and air mixture at a flow rate of 30 mL min^−1^ was used for flame generation. N_2_ at a flow rate of 20 mL min^−1^ was used as the carrier gas for both gas and VFA analysis. The heat of combustion of the solids obtained after lyophilization of starch processing wastewater, before and after dark fermentation, were determined using the bomb calorimeter (Precision Electro Instrumentation India Private Limited, Kolkata, India.

The modified Gompertz equation was used to fit the cumulative hydrogen production obtained in the anaerobic reactors.

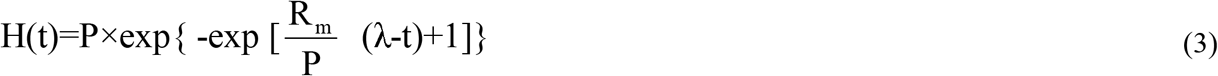

Where H(t) is the cumulative hydrogen production at time t (h), P (mL H_2_) represents hydrogen production potential, R_m_ (mL H_2_ h^−1^) represents the maximum cumulative H_2_ production rate, and λ (h) represents the lag time. The typical cumulative hydrogen production curve was fitted to the above equation [14]. Parameters (P, R_m_, and λ) were estimated using the Curve fitting toolbox ver. 1.1.7 in MATLAB, 2010b.

The performance of the sMFCs was examined in terms of power generation and columbic efficiency. The operating voltage was measured using a digital multimeter with a data acquisition system (USB- 6009, National Instruments, Texas, USA). The anode and cathode potential were measured using a saturated Ag/AgCl reference electrode.

Voltages were recorded every 15 min by a computer (NI LabVIEW-based customized software, Core Technologies, India) and converted to power according to

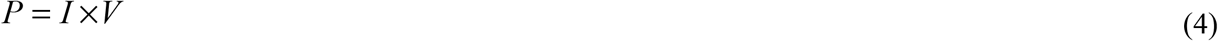

Where *P* = power*, I* = current, and *V* = voltage (V). The volumetric power density was expressed by dividing the power by the working volume of the anode chamber. The current density i_d_ was calculated using

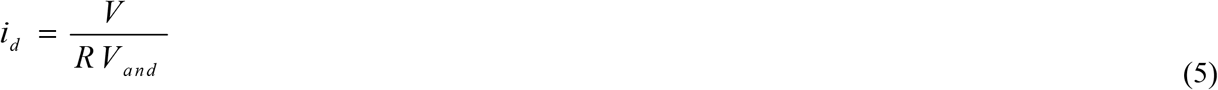

Where *R* is the external resistance (Ω) and *V_and_* (cm^3^) is the working volume of the anode chamber. Polarization curves were obtained using a resistance box (99 kΩ – 0.1 Ω). The Columbic efficiency (CE) is defined as the ratio of total Coulombs transferred to the anode from the substrate to maximum possible Coulombs if all substrate removal produced current. The CE of the MFC operated under batch feed mode over a period of time *t*, was calculated as equation 4 [15]

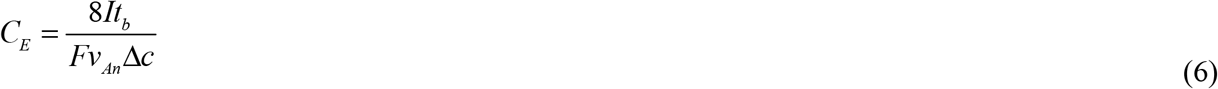

Where *v* is the volume of the anode chamber of MFC.*M* = 32, molecular weight of oxygen; *F*, Faraday’s constant = 96485 C mol^−1^; b = 4, the number of electrons exchanged per mole of oxygen; *q* is the volumetric influent flow rate; Δ*COD* is the difference in the influent and effluent COD.

Short circuit currents (I_sc_) were measured with a Mastech 6000 counts digital multimeter (Precision MASTECH Enterprises Co., Hong Kong) when the anode and cathode were connected directly through the multimeter. The internal resistance of the MFCs was calculated by the current interrupt method [16]. While in closed circuit mode, once the MFC produced a stable current output (I) and potential (V), the circuit was opened causing a steep initial rise in the potential (V_R_) followed by gradual further increment. The steep rise is attributed to ohmic losses caused by the internal resistance (R_int_) of the MFC and can be hence calculated (Eq.7) as

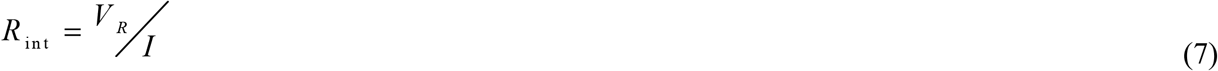

COD removal efficiency can be calculated by the formula

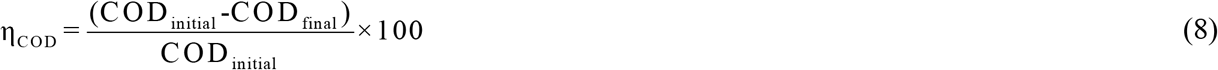

## 3. Results and discussions

### 3.1 Characterization of starchy processing wastewater

Total solids (TS), volatile suspended solids (VSS), pH, COD, total ammonia, and total phosphate were determined and enlisted in Table.1. The waste contained high COD, proving its suitability for the production of H_2_ production process, however, the ammonia content was low and was thus supplemented with an external nitrogen source.

**Table 1.**
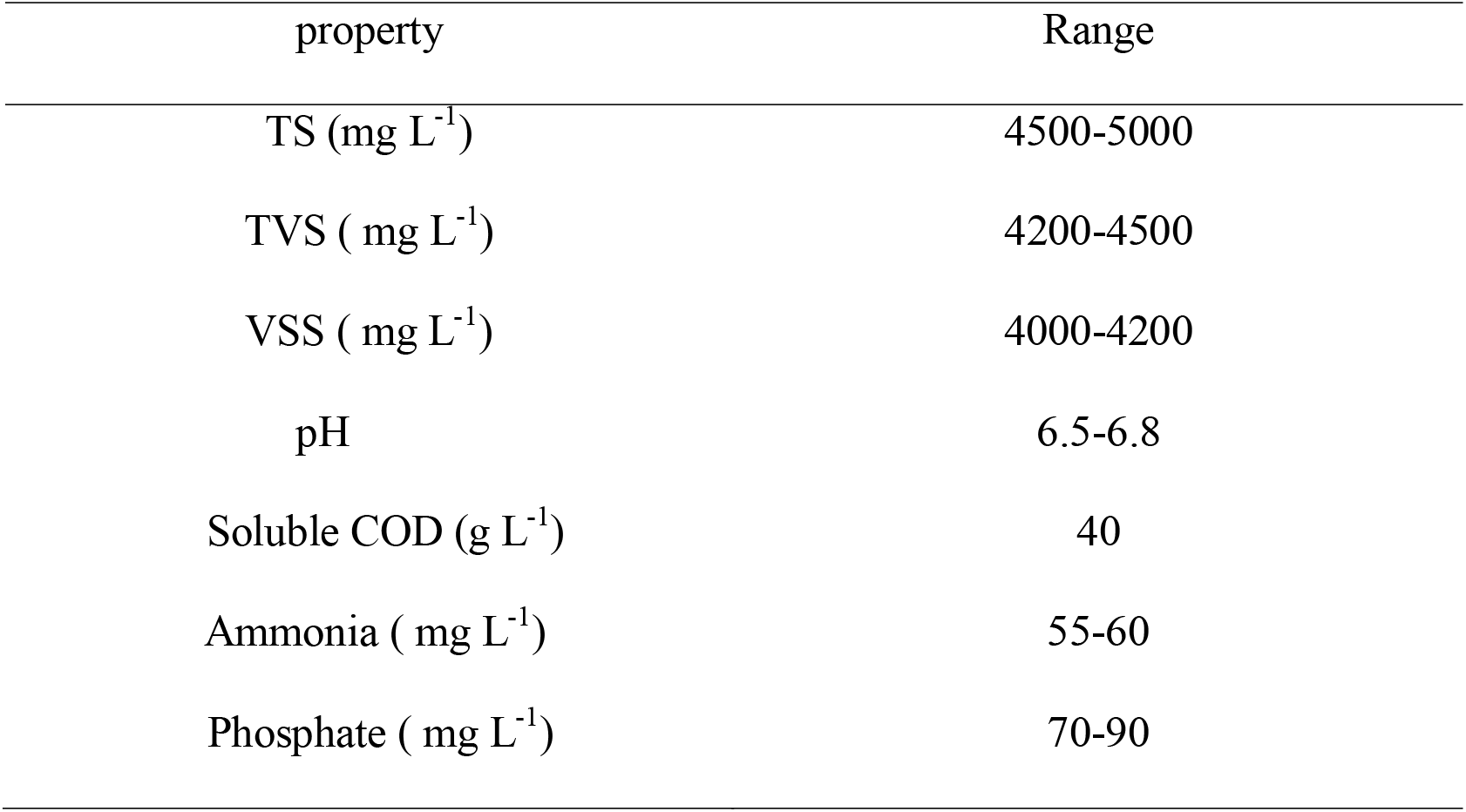
Different physico-chemical characteristics of starchy processing wastewater.

### 3.2 Single parameter optimization

Single parameter optimization studies, with respect to, process parameters (pH, initial COD concentration, and temperature) were achieved using thermophilic enriched mixed culture.

#### 3.2.1 Effect of pH

The effect of initial pH on the cumulative hydrogen production was investigated by varying the initial pH between 4.5-7.5 with an increment of 1 unit. The temperature was maintained 50 ^°^C and initial COD was maintained at 15 g L^−1^. As observed in Fig. 1a, the yield of hydrogen increased from pH 4.5 to 6.5 and thereafter decreased with any further increment in the pH. The maximum cumulative hydrogen production (2.12 L L^−1^) was observed at pH 6.5. The decrement in the hydrogen production at higher pH may be due to the inhibition of hydrogenase activity or due to the redirection of the reducing equivalents towards other pathways [17].

**Fig. 1.**
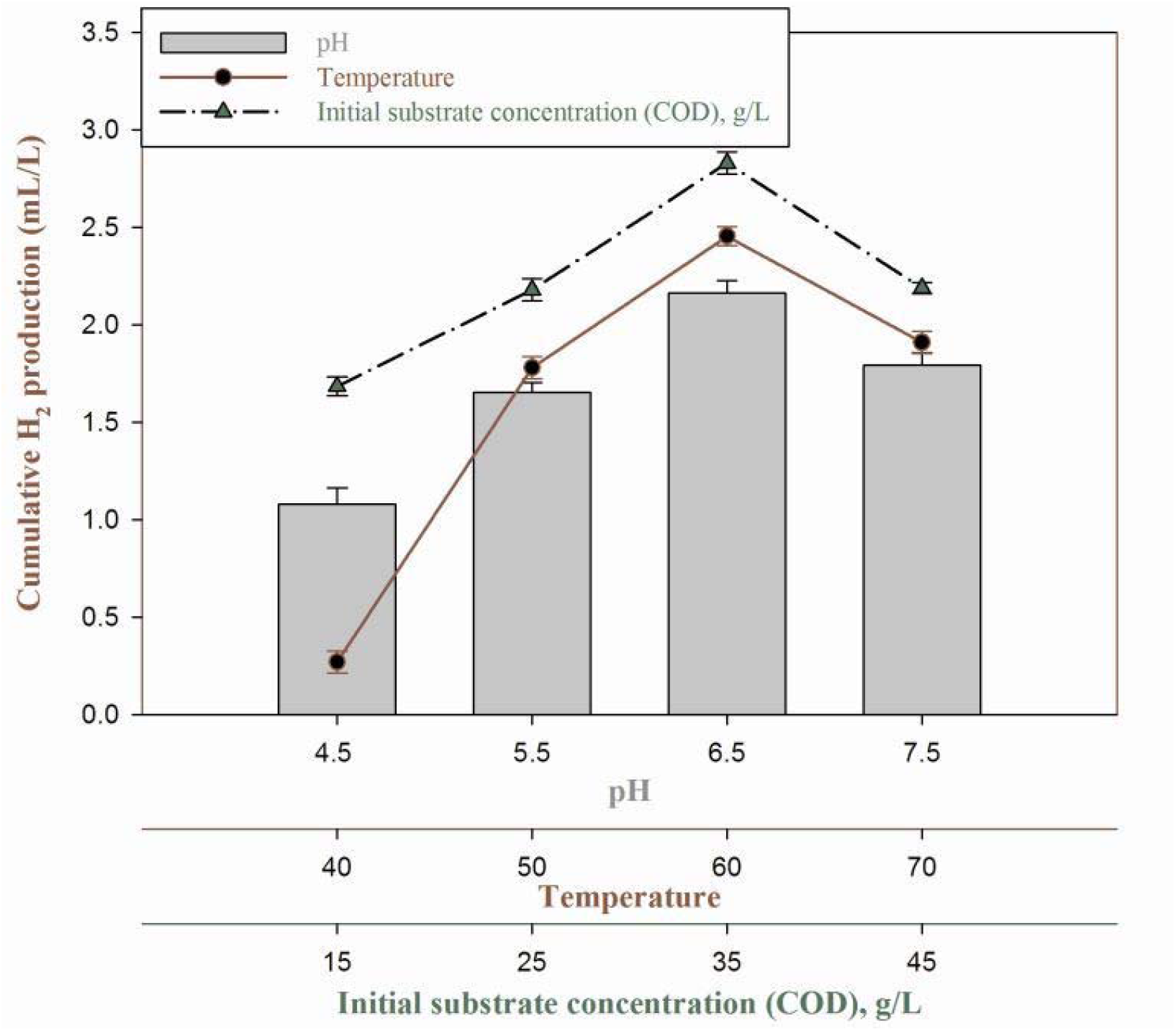
a) Effect of different physico chemical parameters like pH, temperature and Initial substrate concentration on H_2_ production; b) Determination of the activation energy of the mixed culture using Arrhenius equation.

#### 3.2.2 Effect of temperature

Influence of various operational temperatures were investigated at various temperatures from 40 ^°^C to 70 ^°^C at an increment of 10^°^C. The experiments were conducted at initial COD of 15 g L^−1^and optimized pH 6.5. No hydrogen production observed at 40 ^°^C indicating the presence of thermophilic microorganisms in the mixed culture. Hydrogen production increased from 50^°^C to 60^°^C and decreased with further increase in temperature to 70 ^°^C. The reduction of hydrogen production at higher temperatures may be due to the inactivation of the enzymatic machinery [18].

By using Eq.2, the activation energy of the mixed culture was found to be 29 KJ mol^−1^. The value indicated the minimum energy requirement to initiate H_2_ production (Fig. 1b).

#### 3.2.3 Effect of initial substrate concentration

The starch processing wastewater, at different dilutions ranging from 25 % (v/v) to 100 % (v/v) was used in fermentation medium supplemented with the other media constituents, in reactors, set at optimal temperature and pH of 60 ^°^C and 6.5, respectively. Cumulative hydrogen production increased with an increase in initial substrate concentration from 15 g L^−1^ COD to 35 g L^−1^ COD. The maximum cumulative hydrogen production of 2.79 L L^−1^ was observed at 35 g L^−1^ COD. A further increase in the initial substrate concentration to 45 g L^−1^ COD negatively impacted the cumulative hydrogen production probably due to substrate inhibition [19].

### 3.3 Effect of initial substrate concentration on the rate of hydrogen production

Fig. 2 summarizes the batch fermentation of starchy processing wastewater under suitable process parameters. A decrease in pH of the media was observed as the fermentation progressed. The pH drop was in concurrence to the accumulation of volatile fatty acids (VFA) like acetate and butyrate. COD reduction, hydrogen yield and maximum cumulative hydrogen production were 48 %, 6.80 mol H_2_/kg COD _reduced_ and 2.56 L L^−1^ respectively. The soluble metabolite fraction comprised of butyrate (67% w/w), acetate (26% w/w), ethanol (6% w/w) and trace amounts of propionate (1%).

**Fig. 2.**
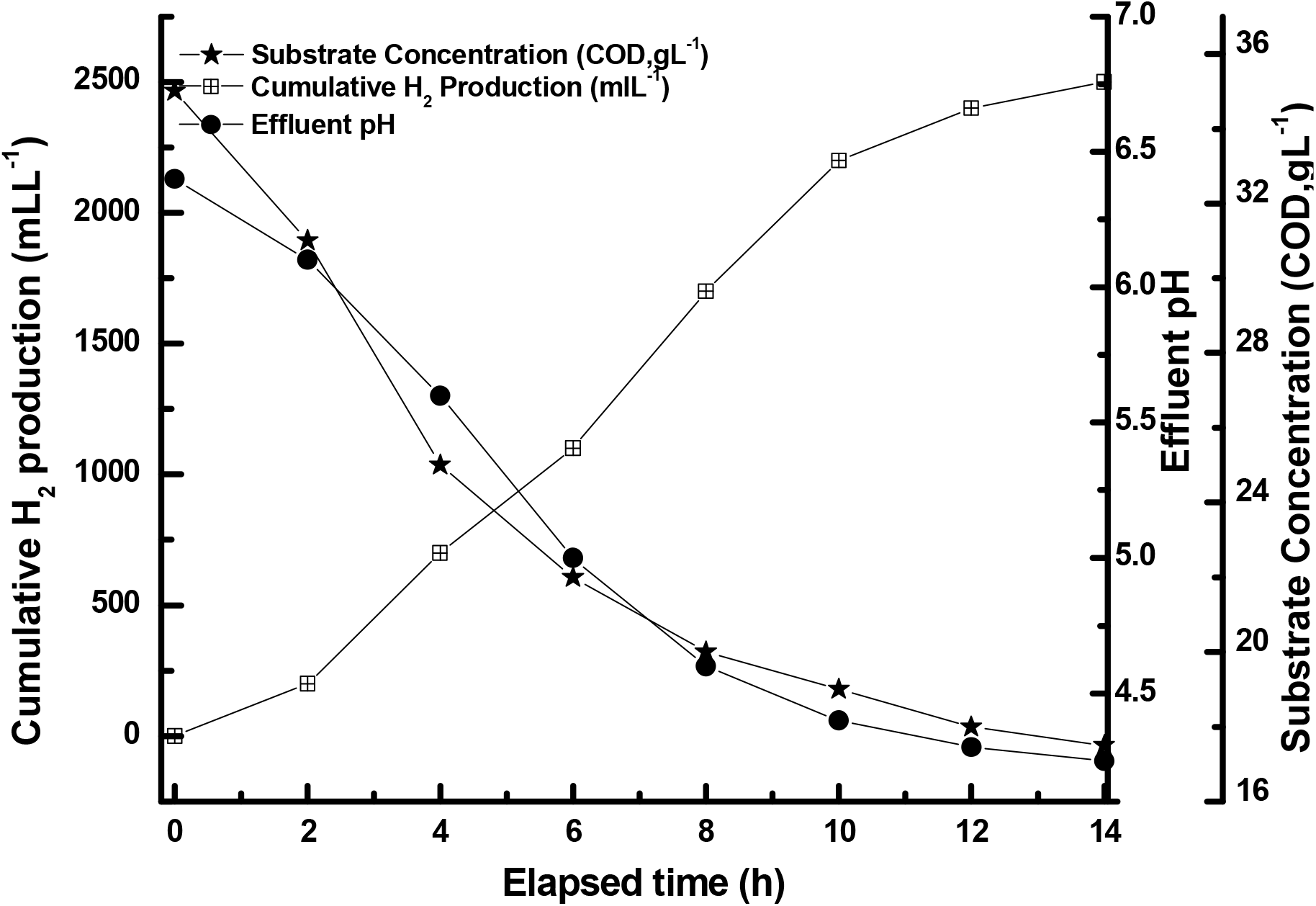
Hydrogen production from starchy processing wastewater at initial substrate concentration COD 35 g L^−1^, initial pH 6.5, temperature 60^°^C.

Different kinetic parameters of hydrogen production process such as lag time (λ), maximum hydrogen production rate (R_m_) and H_2_ production potential (P) at pH 6.5 and temperature 60^°^C were determined using modified Gompertz equation (Table. 2). With an increase in the initial substrate concentration (COD) from 15 g L^−1^ to 35 g L^−1^, lag time decreased. This could be attributed to the availability of greater extent of fermentable sugars corresponding to an increase in the initial COD. However, with further increase in initial COD to 45 g L^−1^, the lag time also increased possibly due to the substrate inhibition at higher substrate concentrations. The H_2_ production potential (P) and hydrogen production rate also followed similar trends with respect to the initial COD.

**Table 2.**
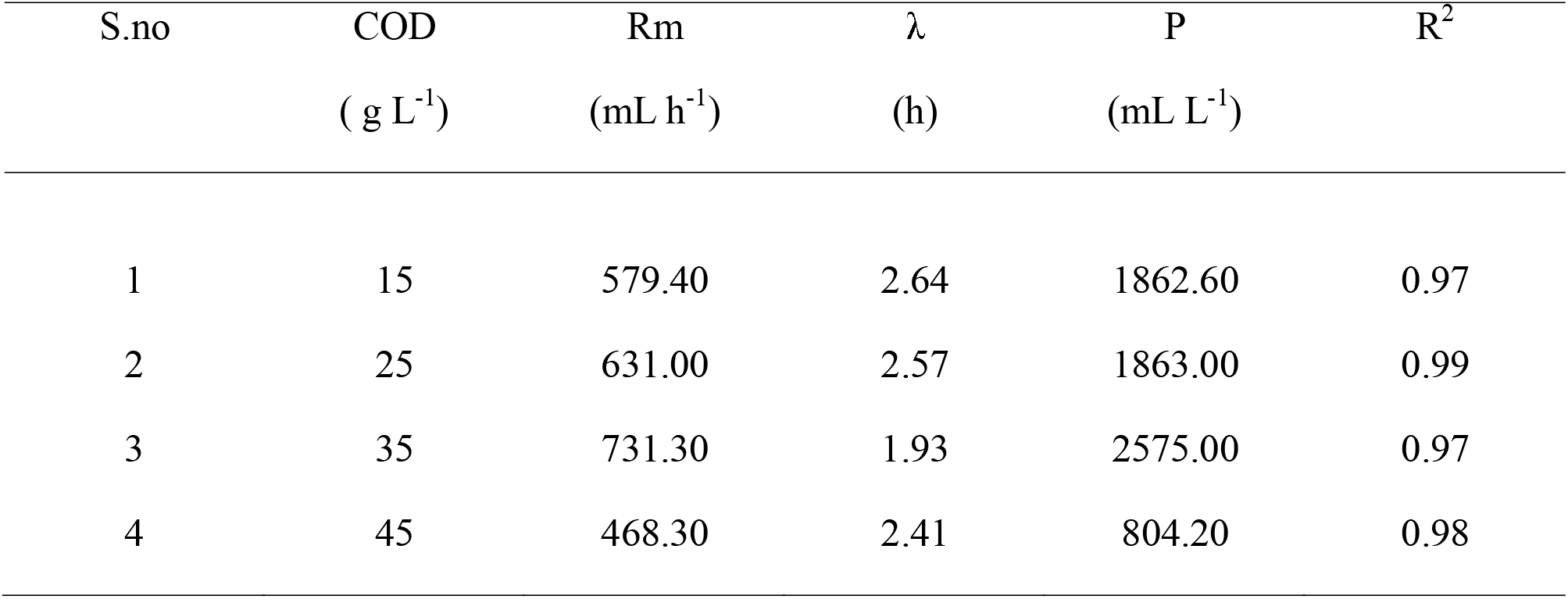
Kinetic parameters of hydrogen production in batch fermentation at different initial COD

### 3.4 Continuous hydrogen production using continuous stirred tank reactor (CSTR)

#### 3.4.1 Effect of organic loading rate

The effect of organic loading rate on hydrogen production, substrate conversion efficiency and biomass concentration (in terms of VSS) in the effluent was studied in a chemostat. A maximum hydrogen production rate of 854 mL L^−1^ h^−1^ was observed at the organic loading rate of 5.60 g L^−1^ h^−1^ (Fig. 3).

**Fig 3.**
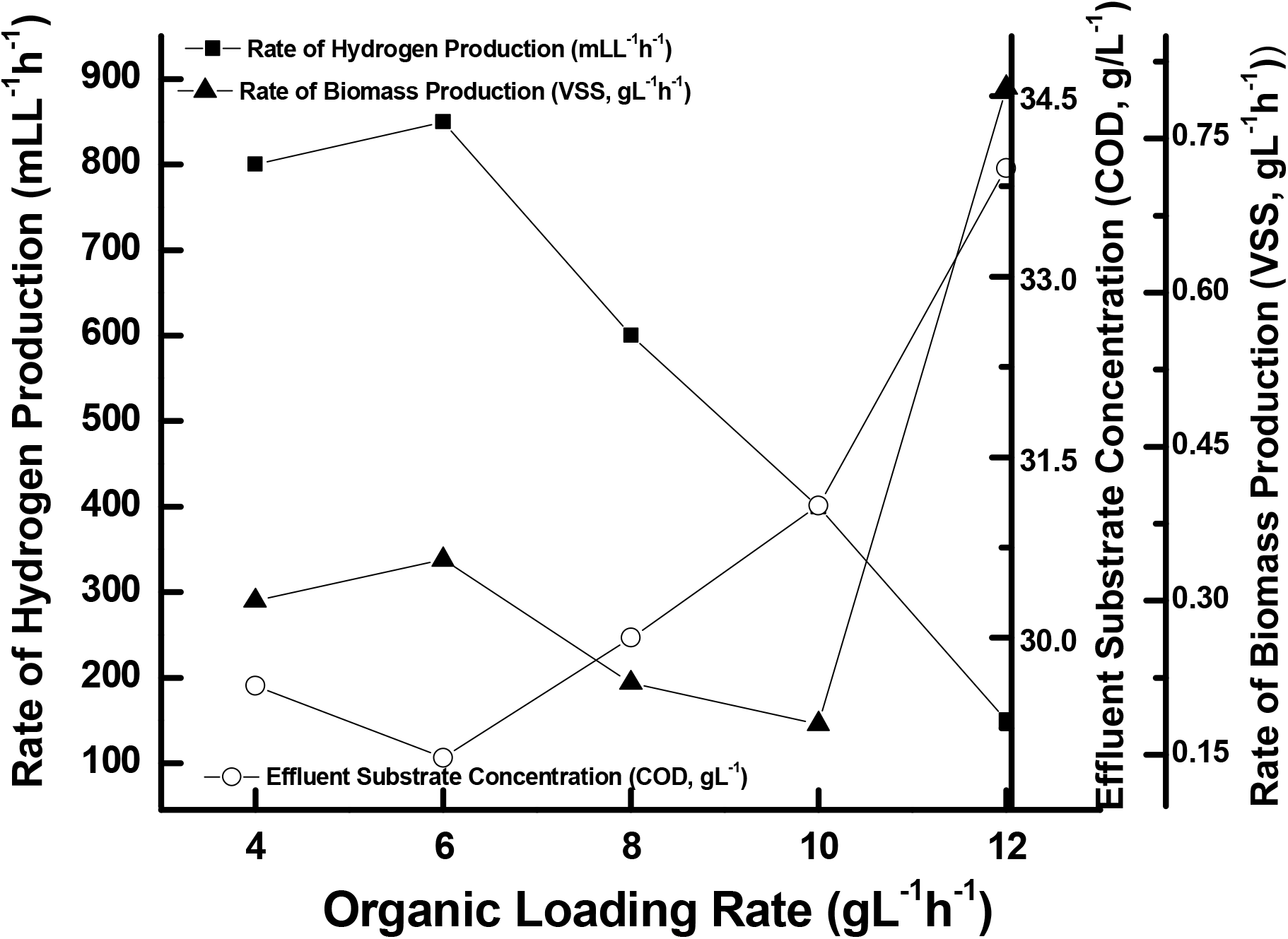
Effect of organic loading rate (OLR) on rate of H_2_ production in CSTR, initial substrate concentration COD 35 g L^−1^, initial pH 6.5, temperature 60^°^C.

The hydrogen production rate was found to be directly proportional with the organic loading rate from 4.20 g L^−1^h^−1^ to 5.60 g L^−1^h^−1^due to the increased substrate availability. With further increase in the OLR, the hydrogen production rate decreased owing to the decrease in the hydraulic retention time (HRT). At higher OLR of 15.05 g L^−1^h^−1^, negligible hydrogen production was observed due to washout of the cells at higher flow rates. Similar studies were carried out to find out the effect of OLR on volatile suspended solids (VSS) and effluent substrate concentration. The biomass concentration and steady state hydrogen production rate followed similar trends. The effect of OLR on H_2_ production rate was reported by several researchers [20,21]. The effluents substrate concentration decreased from 4.20 gL^−1^h^−1^ to 5.60 g L^−1^h^−1^ OLR and then increased. Maximum hydrogen yield of 38.20 mmolL^−1^h^−1^ was observed at OLR 5.60 g L^−1^h^−1^.

The drop in the pH of the effluent was attributed to the accumulation of volatile fatty acids (VFA) formation (Fig. 4 a). The composition of soluble metabolites with OLR and its effect on rate of hydrogen production was studied. The soluble metabolites mainly consisted of butyrate (B), acetate (A) and ethanol. At OLR 5.60 g L^−1^h^−1^, the maximum H_2_ production rate was observed and B/A ratio of the metabolites were found to be 2.46. The effect of B/A ratio on hydrogen production rate was studied by several researchers [10, 22]. In principle, a higher B/A ratio could lead to higher hydrogen production rates due to the availability of ATP which in turn leads to higher biomass formations.

**Fig 4.**
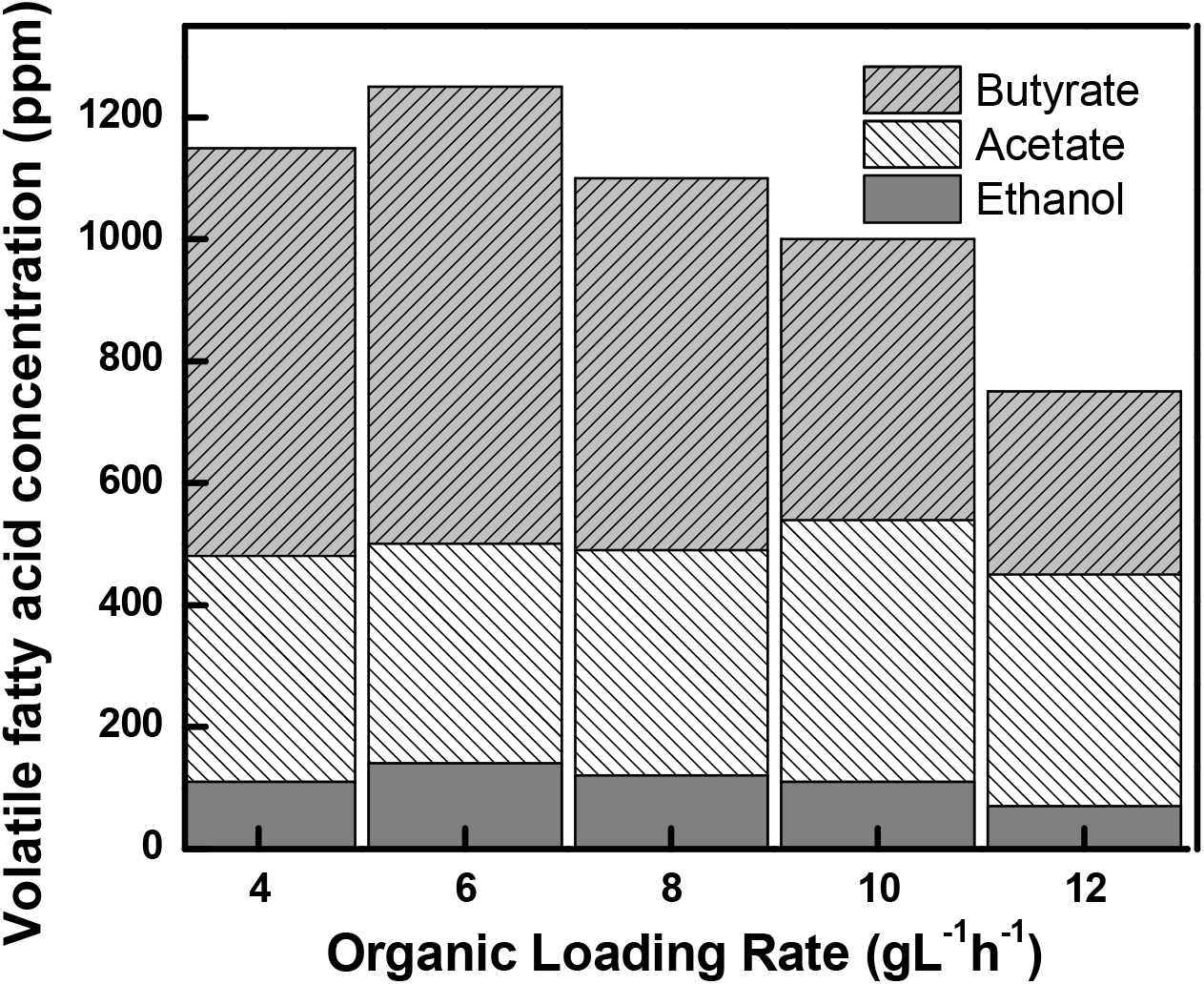
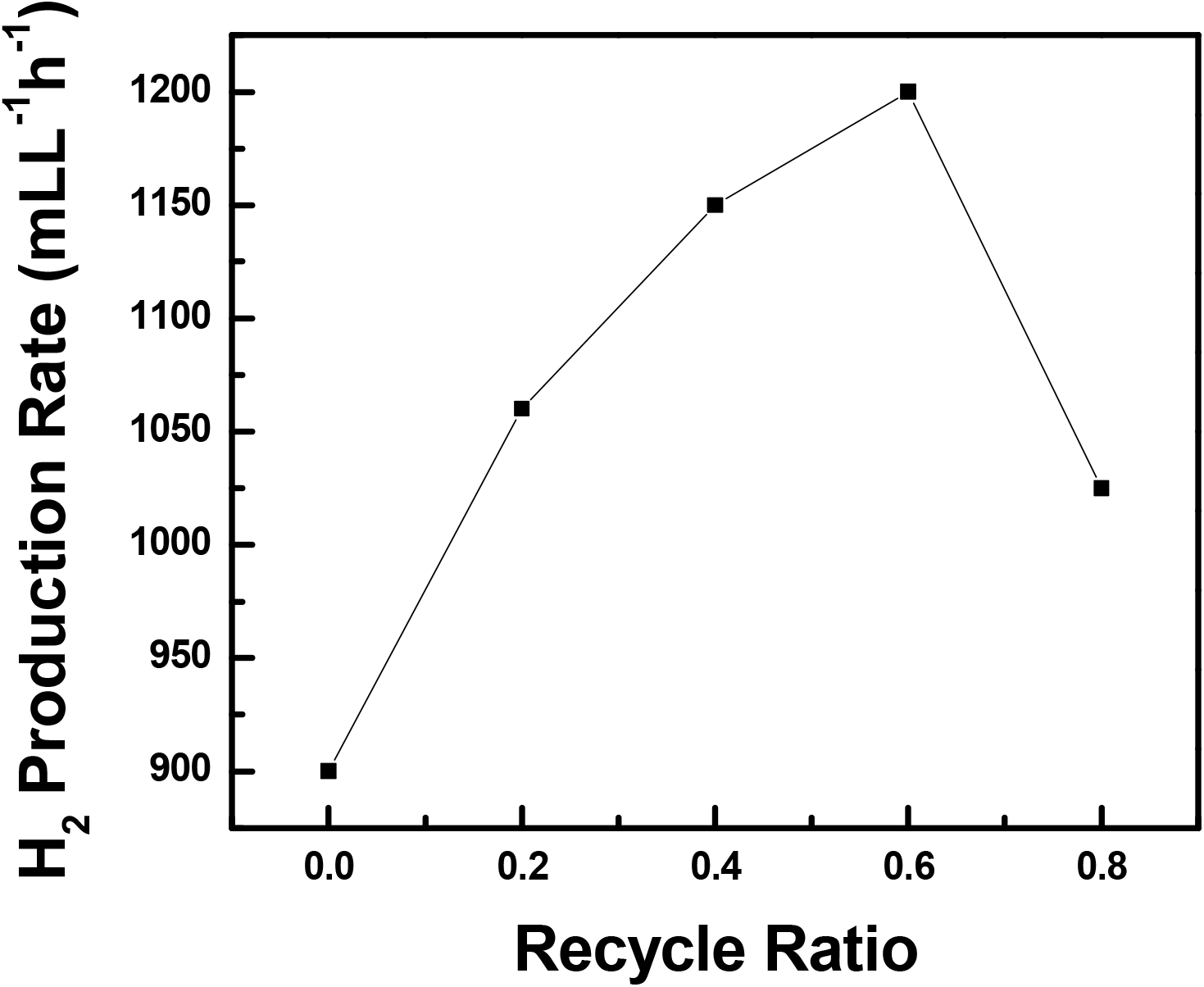
a) Effect of organic loading rate (OLR) on different soluble metabolites and effluent pH; b) Effect of effluent recycle ratio on H_2_ production rate.

#### 3.4.2 Effect of recycle ratio

The effect of effluent recycle on the H_2_ production rate was studied. Recycle ratios were varied from 0 to 0.8 (Fig 4 b). Initially, H_2_ production rate increased with the increase in the recycle ratio up to 0.6 which may be credited to higher solid retention time (SRT) with respect to HRT, which aided in better substrate utilization [23]. A maximum H_2_ production rate of 1224 mL L^−1^ h^−1^ was observed at the recycle ratio of 0.6. Conversely, a further increase in the recycle ratio resulted in a decrease in the H_2_ production rate probably due to an increase in the substrate concentration in the reactor that led to substrate inhibition. Moreover higher recycle ratios may have led to a drastic drop in the pH in the media owing to a higher inflow of VFAs into the system.

### 3.5 Electricity generation using MFC

Spent medium collected from the batches of dark fermentation was used to study its suitability as anolyte in the MFCs for further energy recovery. Prior to use, the effluent was centrifuged at 5000 rpm to remove the cell suspension from the spent media followed by neutralization of the wastewater, since the spent media of starchy wastewater is acidic, as deemed necessary for further biological treatment. Higher acidity or alkalinity of wastewater affects both wastewater treatment efficiency and the environment inside the reactor. Since the COD of the spent media was around 18.2 g L^−1^; it was diluted to around 4.5 g L^−1^ and 100 mL of that was used as anolyte for each cycle. The pH of the spent medium was adjusted near to 7.5 for electrogenesis in MFC-1 and 2 through a buffer (carbonate) and alkali supplementation respectively; contrariwise, the pH of the anolyte in MFC-3 remained unaltered (nearly 5). Six days after inoculation, the anode half cell potential decreased owing to the donation of electrons to the anode by electrogenic bacteria; and the voltage reached a plateau of about −160 ± 6 mV and 143 ± 6 mV (vs. Ag/AgCl) against an external resistance of 100 Ω for MFC-1 and MFC-2 respectively,. However, significant variation was found in the anode half-cell potential of MFC-3. The maximum anode potential reached −80 mV vs. Ag/AgCl (100 Ω); which may be attributed to the fact that presence of higher acidic conditions in anolyte hampers electrogenic activity of the EABs. Both sMFC-1 and 2 took a period of around 11 days to reach stable conditions under batch mode of operation, at feed cycle time of 36 h. Slow increase in the current was observed during the duration of the operation. Given an external load of 100 Ω, the current production reached the maximum value of 1.67, 1.44 and 0.8 mA for MFC-1, 2 and 3 respectively on the 11^th^ day of operation using the diluted spent medium. At the same time, a maximum volumetric power density of 4.19 Wm^−3^ and 3.7 Wm^−3^ was generated with MFC-1 and MFC-2 respectively. The power densities obtained showed that MFC- 1 produced 12 % higher volumetric power densities as compared to MFC-2; these results clearly indicated that the spent media neutralized with carbonate buffer as a substrate had greater potential for higher power generation. During the polarization, the anode potentials observed in all the three sMFCs were found to be different which indicated that the effect of the microbial activities on the anode was responsible for the voltage losses and the substantial limitations were from the anode side. The corresponding anodic half cell polarization curves of the MFCs (Fig. 5) were obtained by varying the external resistance from 90 tí! to 40 Ω. The final anolyte pH of MFC-1 and MFC-2 were documented as 6.82 and 5.85 respectively at the end of the 11^th^ batch cycle. The result suggested that the inferior performance of MFC-2 could be attributed to the pH imbalance associated with voltage losses: the accumulation of proton in highly saline anolyte (during neutralization with alkali addition conductivity was found 16.9 mS cm^−1^) and transmembrane transport of the other cations created due to pH splitting. The protons produced during the microbial catabolism in the anode chamber were not consumed and rather resulted in pH reduction in the anode chamber causing a decrease in the MFC performance. In MFC-1, carbonate buffer addition facilitied in maintaining the anolyte pH close to the optimal value.

**Fig 5.**
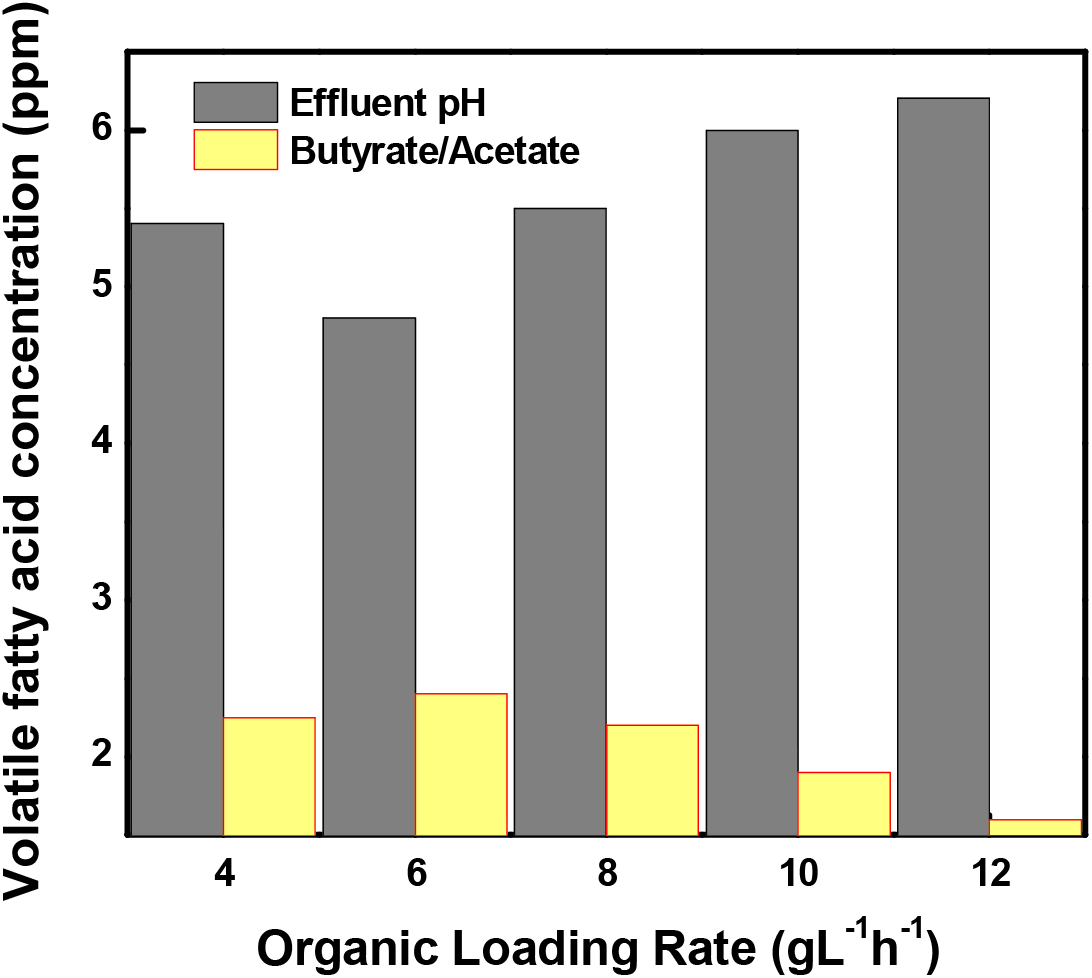
Polarization study using different anolytes: buffer treated (MFC-1), alkali treated (MFC-2) and without treatment (MFC-3).

### 3.6 Sustainable power

Relative deviation in the anodic potential (RDAP) with the function of applied external resistance was used to evaluate the maximum sustainable power which was calculated using the initial anodic potential (E_o,anodic_) and the anodic potentials generated at applied external resistance. Power generated by MFCs can be computed as the product of the cell potential and the current in the external circuitry [24]. The fuel cell and the external circuit will be in steady state if the power generated by the MFC equals the power consumption for an extended time and at steady state conditions the power production is sustainable. Because many steady states are possible, it is important to define conditions at which the sustainable current reaches a maximum by a MFC [25]. A plot between the variation in percent deviation of anodic oxidation potential (RDAP) with the function of applied external resistance was used to evaluate the external resistor to measure the maximum sustainable power of MFC.

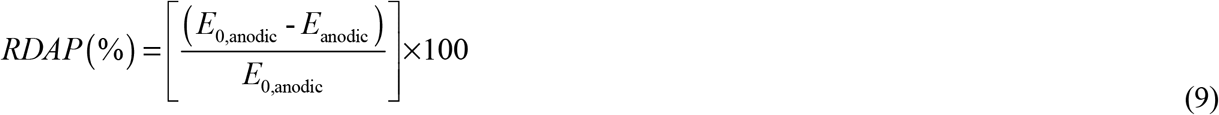

The percent variation of RDAP with respect to applied external resistance is shown in Fig. 6 a & 6 b. The linear fit at higher applied external resistance represented a region in which the external resistance controled the power, while, the linear fit at lower applied external resistance represented a region in which the power was limited by kinetics, mass transfer, or internal resistance (Menicucci et al., 2006). The performance of the fuel cell was considered to be in a steady state if the power generated by the system equaled the power consumed over an extended time. Moreover, at steady state power production is considered to be sustainable. When external resistance was high, the RDAP increased linearly with decreasing external resistance because the electron delivery to the cathode was limited by the external resistance. At lower applied external resistance, the electron delivery to the cathode was limited by kinetic and/or mass transfer (or internal resistance) and the RDAP increased linearly with decreasing external resistance. The conditions where external and internal resistances were equal between these two lines, a horizontal line from the intersection was drawn to estimate the external resistor that allowed measuring the sustainable power. RDAP analysis documented lower sustainable resistance with MFC-1 (7 kΩ) as compared to MFC-2 (8 kΩ) and MFC-3 (14 kΩ) indicating a decrease in overpotential and an increase in system sustainability with the addition of carbonate buffer in anolyte (Fig. 6). The corresponding sustainable power densities of the three MFCs were 0.5 Wm^−3^; 0.4 Wm^−3^ and 0.29 Wm^−3^ respectively. This result suggest that the reduction in anolyte pH may have detrimental effect on the metabolic activity of EABs which impeded the electron donations in anode, as a result of the power found to a lesser extent in MFC-2 as compared to MFC-1.

Fig. 6a. shows the effect of external resistance on the variation of the percent deviation of anodic potential with respect to the applied external resistance and evaluating the sustainable power in sMFC, A. MFC 1; B. MFC 2.

### 3.7 Bioelectrochemical characterization

Electrochemical Impedance Spectroscopy (EIS) can be performed in a three electrode cell (with working, counter, and reference electrodes) to investigate the electrochemical processes for the individual electrode, or in a whole cell configuration (only working and counter electrodes). To calculate the impedance, data was analyzed by equivalent circuit (EC) modeling. Common impedance components are resistance (R), capacitance (C), and inductance (L). Internal resistance (R_int_) mainly consists of three parts: charge transfer/polarization/activation resistance (R_ct_), solution resistance (R_S_), and concentration/diffusion resistance named as Warburg resistance (W). R_ct_ value directly related to the electron transfer behavior of an electrochemical reaction occurring in the electrode. A significant difference in the semicircle region has been observed in the impedance plot of the electrodes (Fig. 7). The R_ct_ value of the different MFCs trailed the following order: MFC-1 (buffer pretreatment) (97.4 Ω) < MFC-2 (132.09 Ω) < sMFC-P (168.86 Ω). The minimum R_t_ value obtained with buffer pretreated MFC indicated the maximum electron transport due to the oxidation of the acid rich spent media which reduced the anodic voltage losses and increased the current output. The results obtained from EIS support the results obtained from the half cell polarization study. The internal resistance in MFC-3 originated from the poor biofilm development on the anode surface and subsequent reduced electrochemical reactions on the anodic surface.

### 3.8 Wastewater treatment

The wastewater treatment performance of all the MFCs was observed in terms of COD removal at the different operating cycles. The results showed that the average COD removal efficiency of MFC-1 and MFC-2 was in the range of 72-80 %, which demonstrated the effective treatment efficiency of this system. Coulombic efficiency (CE) is the key parameter used to evaluate the recovery of the electron through the external circuit against that theoretically present in the organic matter of the MFCs. After 2 weeks, the average CE observed in MFC-1 and MFC-2 was 8.2 % and 6.8 % respectively (Fig. 6b & Fig. 6c). MFC-1 showed higher value owing to better metabolic activity of EAB in the reactor as compared to MFC-2. The Columbic efficiency of the MFCs were very less indicating there was a substantial COD that was not associated with power generation. MFC-1 showed higher columbic efficiency during all the operating cycles, however, CE was slightly improved in MFC-2 at a later stage probably due to the development of a bio-film on the anode, owing to reduction in the methanogens in the inoculum at slightly lower pH of nearly 6.

The volatile fatty acids (VFA) produced in the acidogenic fermentation was found suitable as substrate for bioelectricity generation. The reduction of COD and total carbohydrate of wastewater were 73 % and 58 % respectively. This study demonstrate that additional renewable bio-energy with simultaneous wastewater treatment can be achieved by utilizing acid-rich effluents generated from acidogenic fermentation without any external power consumption.

### 3.9 Energy recovery calculation from two stage system

## Energy efficiency in Dark fermentation

Heat of combustion value of H_2_ = 12.64 KJ L^−1^

Heat of combustion of starchy processing wastewater = 22.4 KJ g^−1^solids.

Total solids obtained from liter of the medium = 15 g.

Total heat of combustion of the fermentation medium used in dark fermentation = 336 KJ

Volumetric yield of H_2_ from batch fermentation under suitable conditions = 2.56 L L^−1^

Total Heating value H_2_ evolved from batch fermentation = 2.56 L L^−1^ * 12.64 KJ L^−1^

= 32.36 KJ L^−1^.

Energy recovery from feed = (Heat of combustion of H_2_) *100/Heat of combustion of feed.

= 32.36*100/336

= 9.63%.

### Energy efficiency in MFC

Power output in MFC was calculated as P (Watt) = I × V and Energy production can be expressed as

E (Joule) = P × t

(8)

The start up of the power production, expressed as Wm^−2^ electrode surface, stable power generation was achieved after 4^th^ cycle onwards. In each cycle the sMFC was dosed with anolyte containing buffer or alkali treated diluted spent media (3000 mg L^−1^).

1000 ml contain COD = 3000 mg

So, 100 ml contain COD = (3000/1000) X100 mg=300 mg.

COD removal = 73 % (after 36 h)

COD utilized = 219 mg.

Heat of combustion of 1000 mg = 14.72 KJ

So, 219 mg = (14.72* 219)/1000 = 3223.68 J.

For the dose of 3 g COD L^−1^ to the 100 mL liquid in the anode compartment, representing 3223.68 J available energy, the average energy production as electricity n MFC-1 (per batch cycle) according to Equation (8) with 0.288 W m^−2^ × 0.0016 m^2^× 36 h × 3600 s h^−1^ = 59.72 J.

Energy efficiency for a unit MFC-1 per batch cycle = (59.72/3223.68)*100=1.86 %

Hence, an energy recovery as electricity of 1.86 % was achieved through a unit MFC.

Several reports of two stage processes are available where the dark fermentation is coupled with MFC. However, the present process showed higher rate of hydrogen production (1224 mL L^−1^ h^−1^) and improved power density as compared to all the reported systems. Moreover the COD reduction of the present system is appreciable when compared to the reported systems (Table 3).

## 5 Conclusion

An enriched thermophilic mixed culture was found suitable for the treatment of starchy processing wastewater. The spent media of dark fermentation was utilized for the power generation and COD reduction. The phosphate concentration decreased slightly during the dark fermentation process. But in the bioelectrochemical system the concentration of phosphate decreased significantly. This reduced the risk of eutrophication when discharged in the environment. Thus the established the two stage process offered a higher energy recovery from the wastewater coupled with a lower risk of environmental pollution when discharged directly into the environment.

## Acknowledgement

The authors wish to thank the financial support from MNRE, CSIR, UGC, and DBT, Govt. of India and IIT Kharagpur for the facilities.

